## Supplement Figs and Table 1, 3 and 4 for "Coordinated conformational changes in the Tad pilus ATPase CpaF facilitate a rotary mechanism of catalysis"

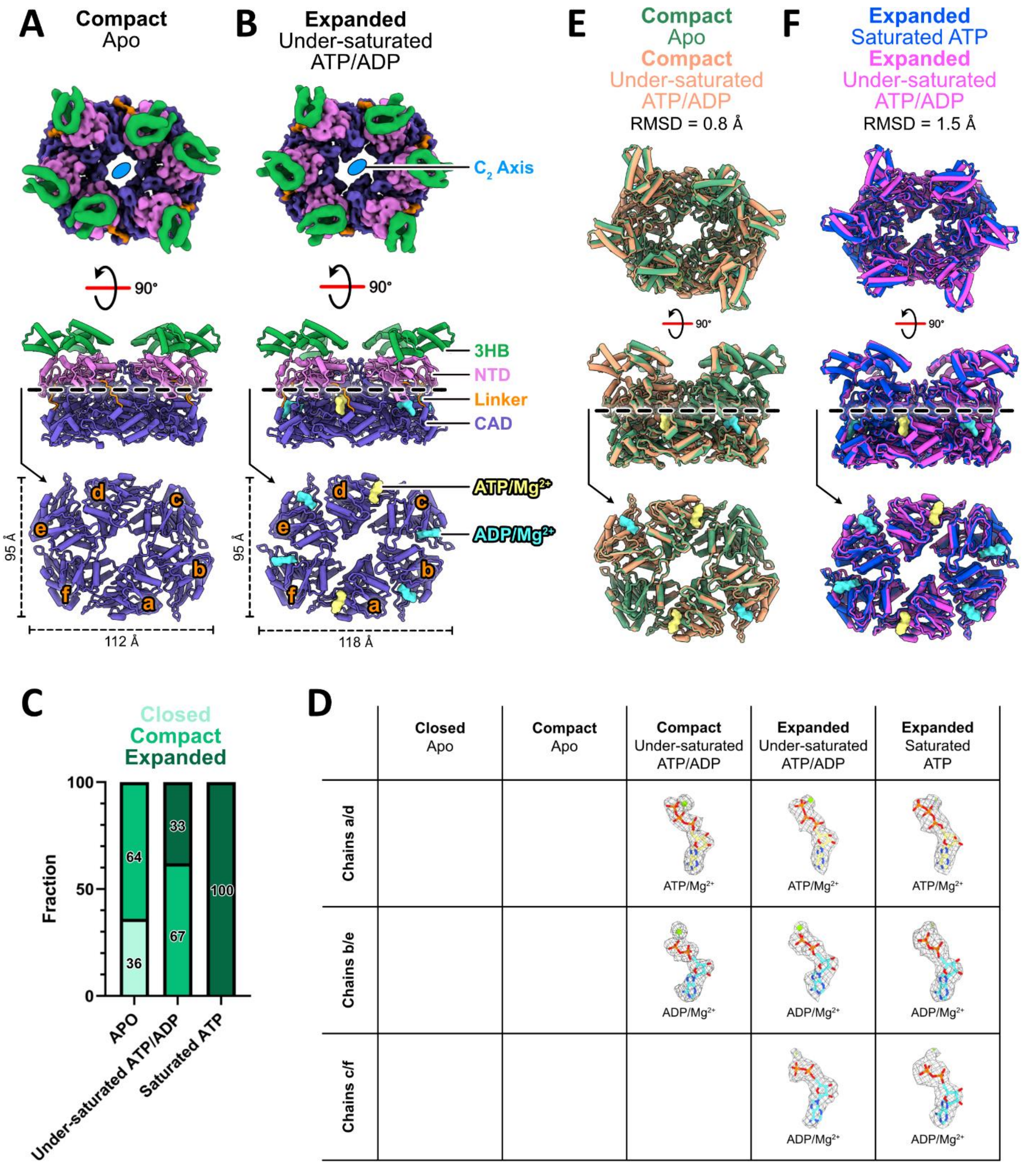


**Supplementary Figure 1. Additional CpaF structures, particle distributions, nucleotide densities, and structural alignments. A-B.** Additional CpaF structures in the compact and expanded conformations determined from the apo and under-saturated ATP/ADP datasets, respectively, represented as described in Figure 1. **C.** Distribution of particles that contributed to the reconstruction of each conformation for the three datasets. Increasing nucleotide concentrations favor the expanded state. **D.** Cryo-EM densities in the active site pockets of all five structures obtained from the three datasets, and the nucleotides modeled into the density. All cryo-

EM densities within each dataset are depicted at the same threshold. **E-F.** The RMSD of the two aligned compact structures from the apo and under-saturated ATP/ADP datasets, and the two expanded structures from the under-saturated ATP/ADP and saturated ATP datasets, was calculated along the entire hexamer.

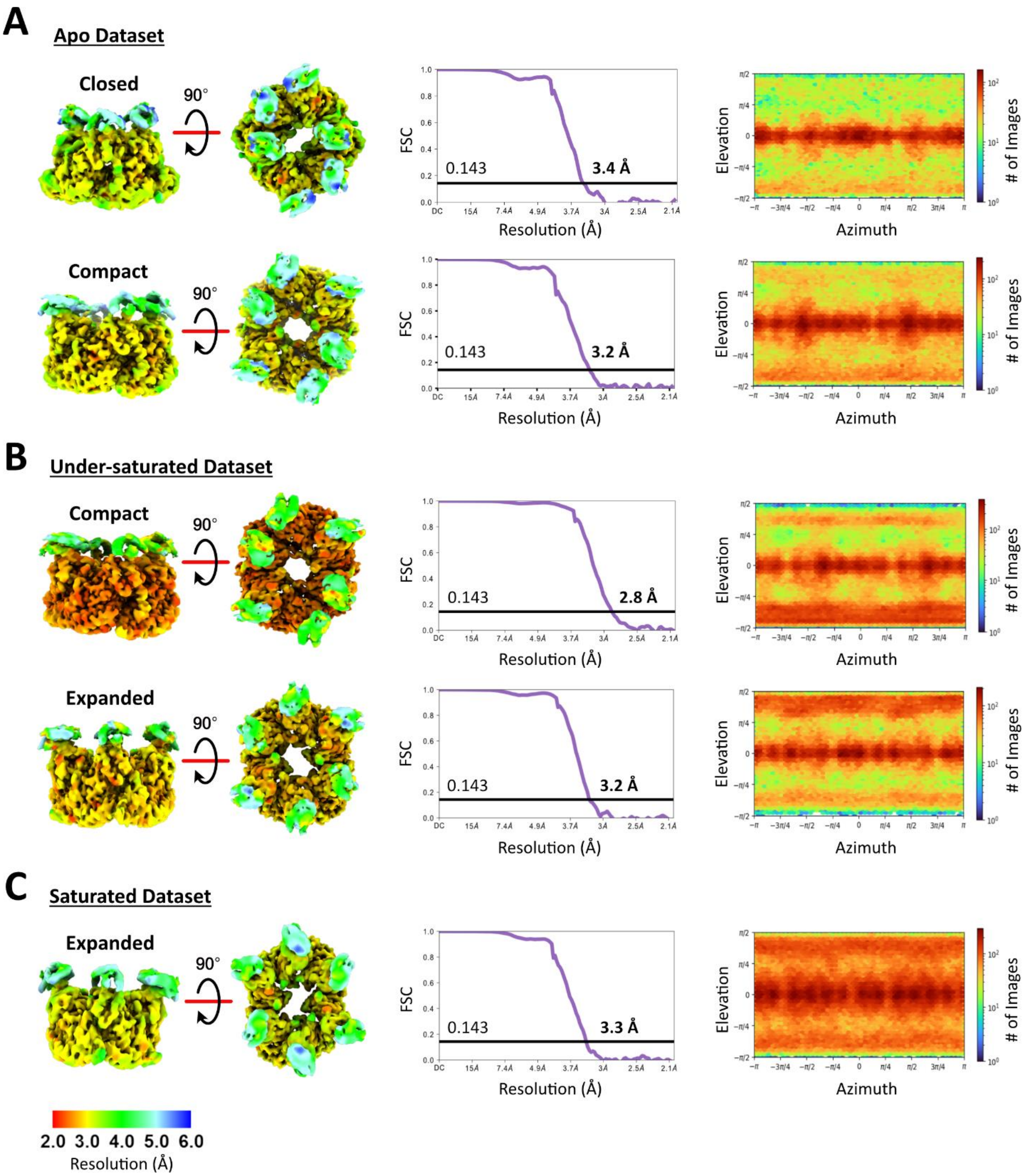


**Supplementary Figure 2. Consensus cryo-EM map validation. A-C.** (left) Consensus maps of the biological hexamer from the three datasets, depicted in top and side views and colored by local resolution, along with the (middle) corrected Fourier shell correlation (FSC) curves following a gold standard refinement and the (right) orientation distribution plots.

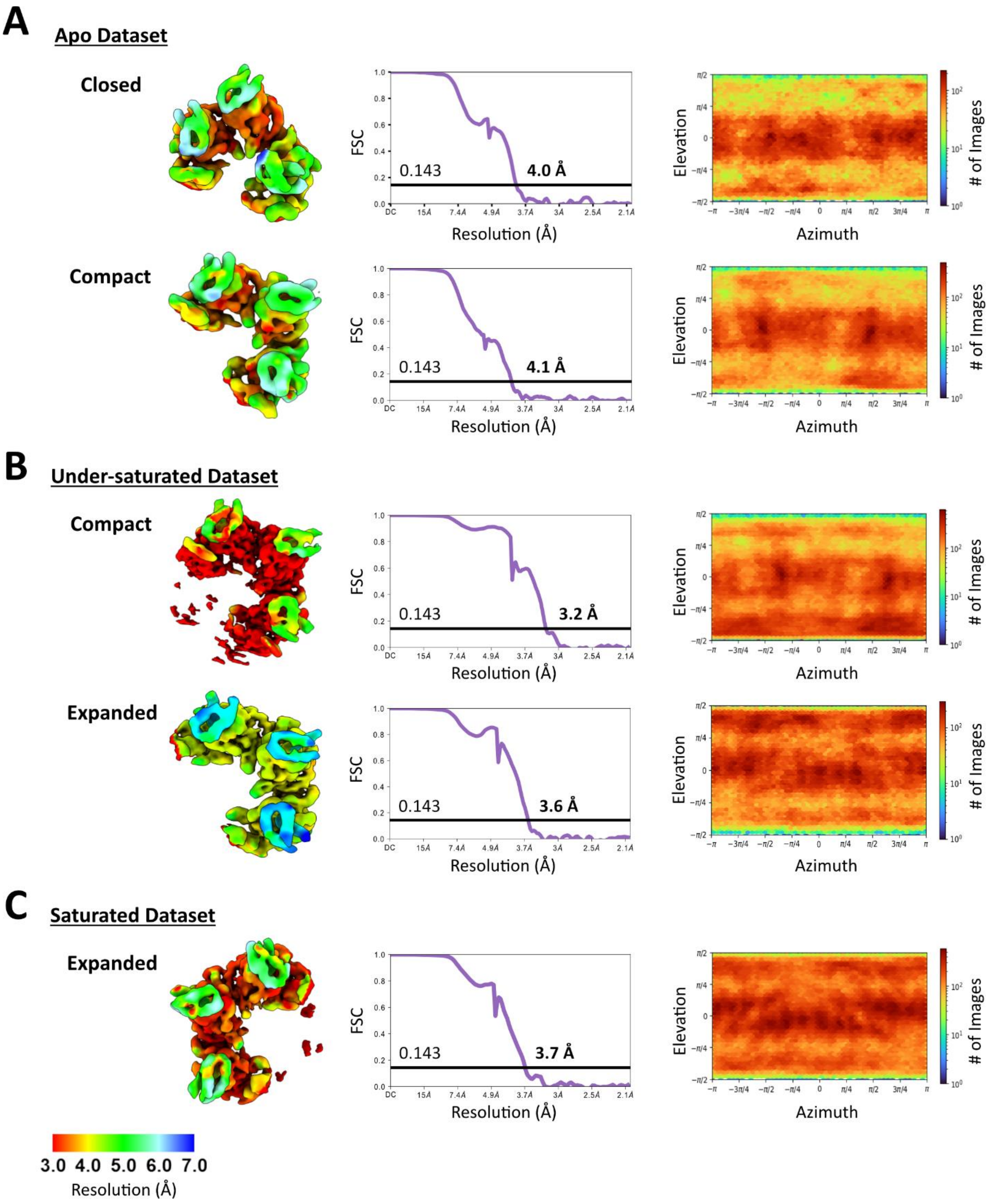


**Supplementary Figure 3. Local refined cryo-EM map validation. A-C.** (left) Locally refined maps of the 3HBs and NTDs from the asymmetric trimer in the three datasets, depicted in top view and colored by local resolution, along with the (middle) corrected FSC curves following a gold standard refinement and the (right) orientation distribution plots.

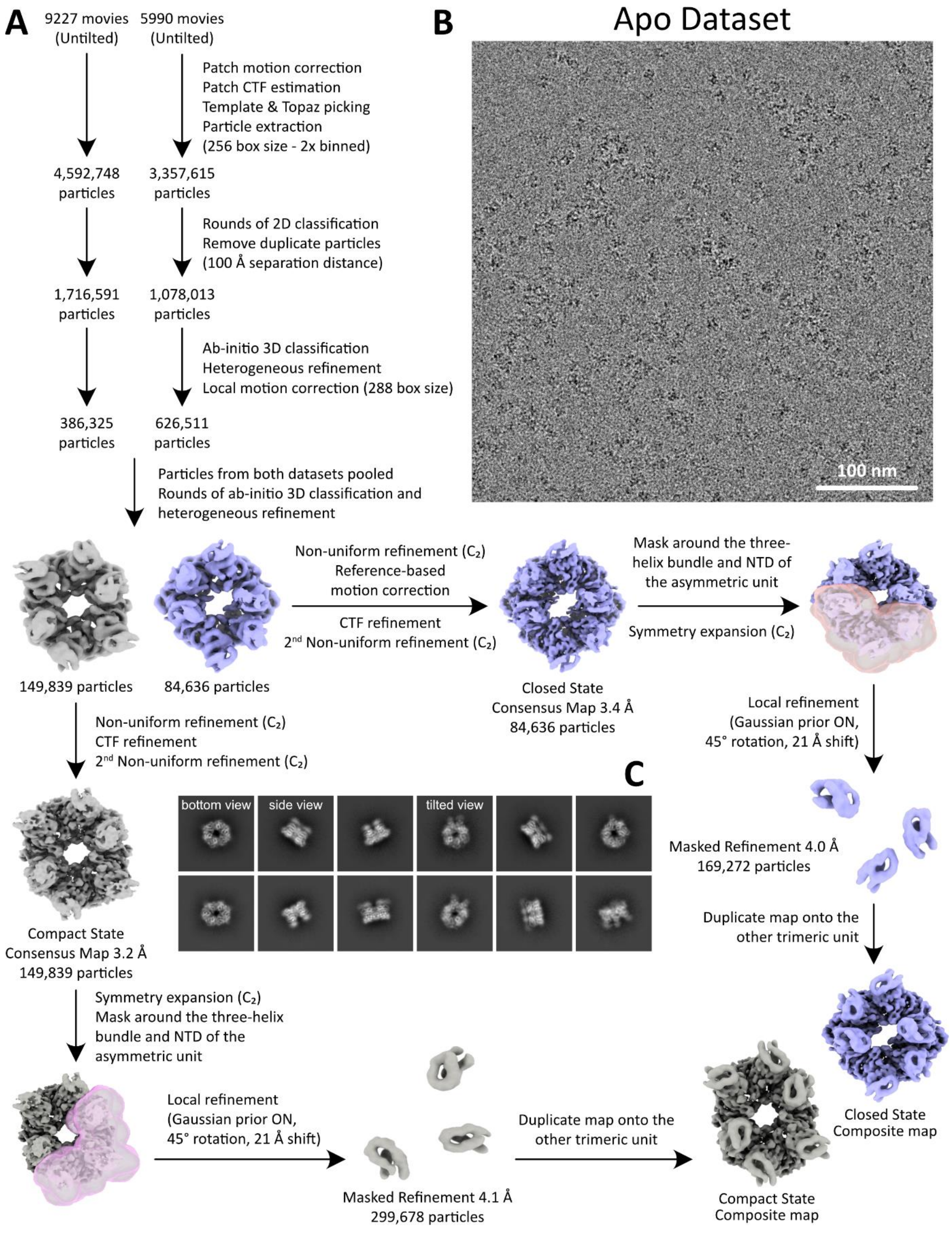


**Supplementary Figure 4. Image processing workflow for the apo dataset. A.** cryoSPARC processing workflow for obtaining consensus, local refined, and composite maps of the closed and compact states. **B.** Representative micrograph and **C.** 2D class average images of both states.

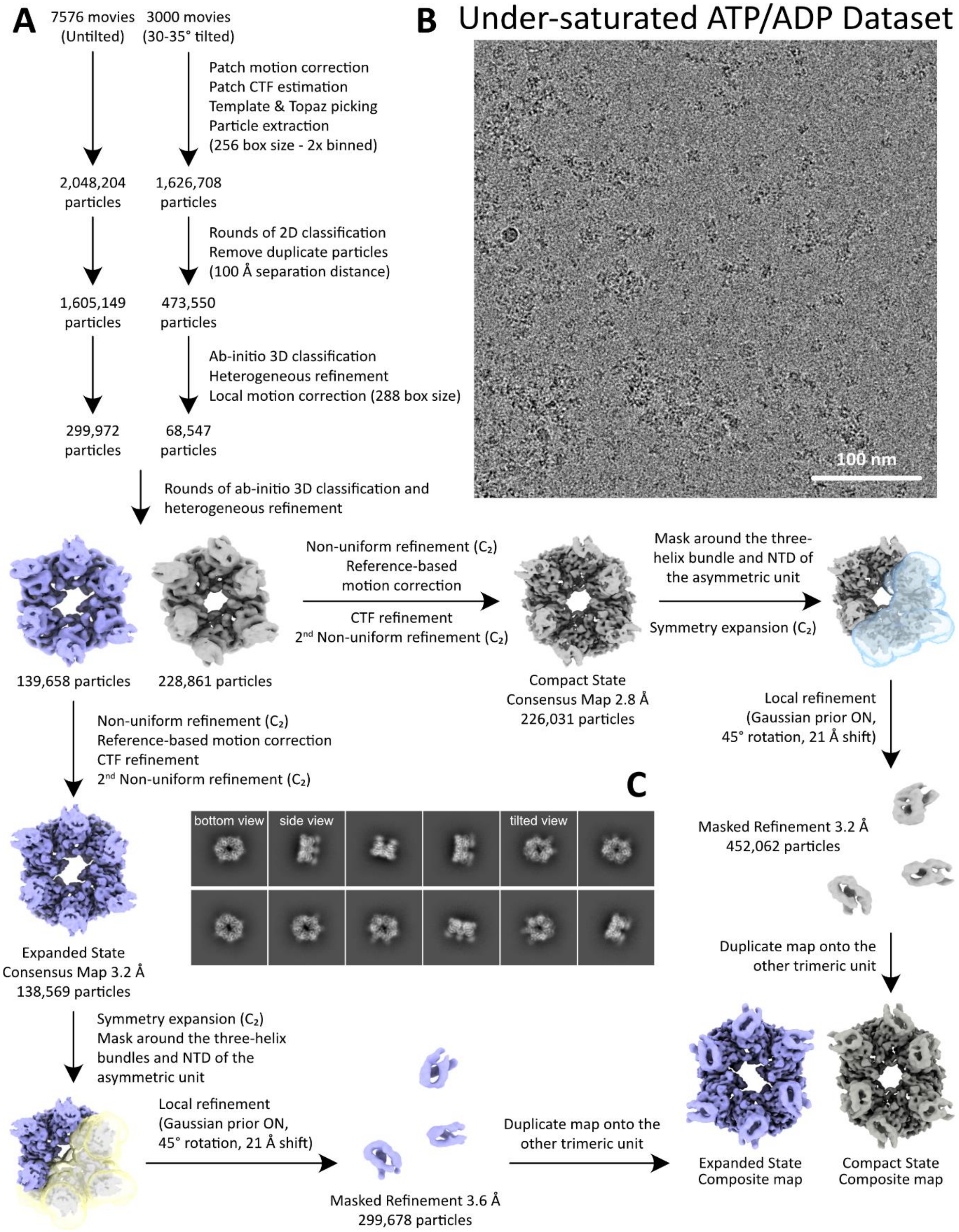


**Supplementary Figure 5. Image processing workflow for the under-saturated ATP/ADP dataset. A.** cryoSPARC processing workflow for obtaining consensus, local refined, and composite maps of the compact and expanded states. **B.** Representative micrograph and **C.** 2D class average images of both states.

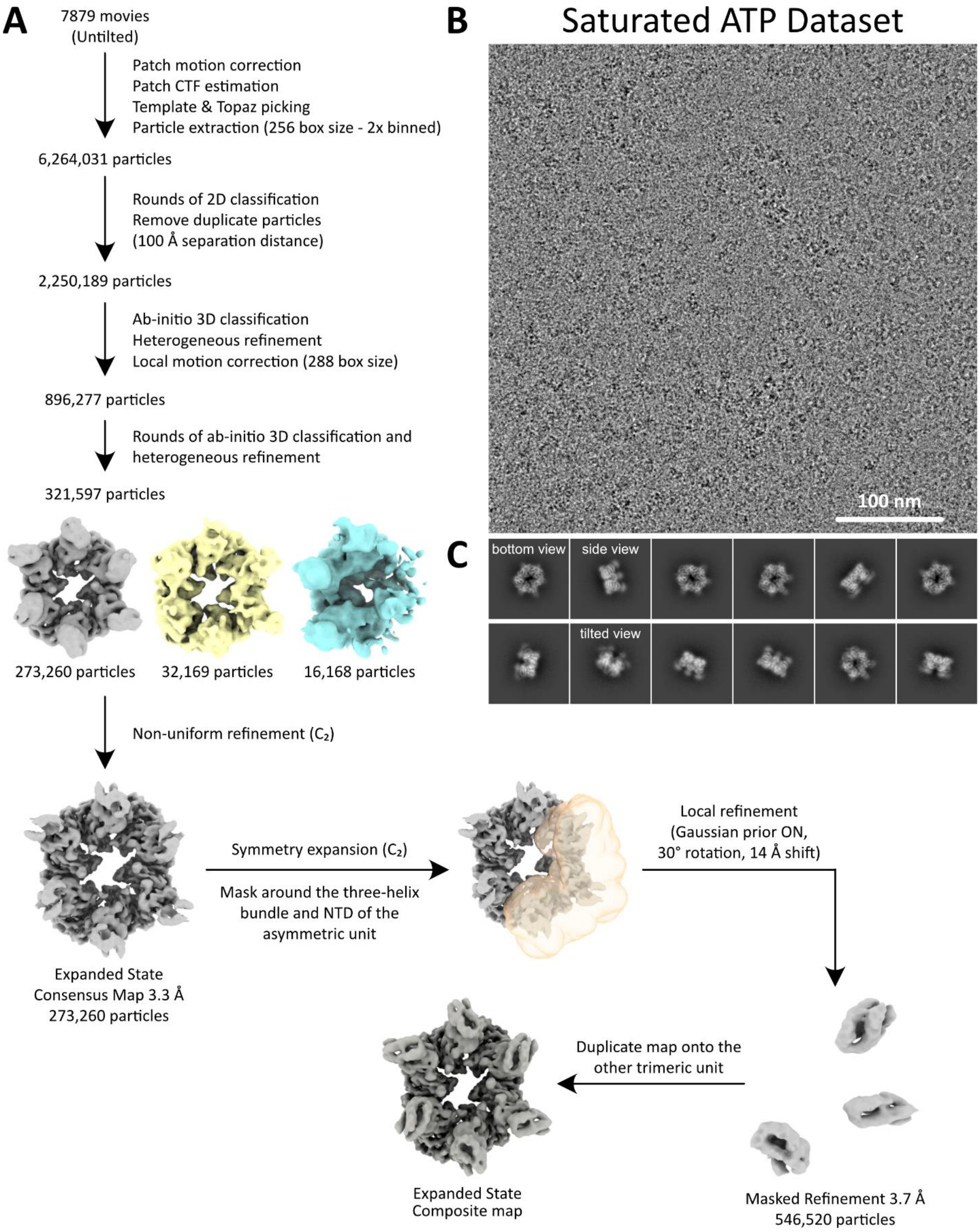


**Supplementary Figure 6. Image processing workflow for the saturated ATP dataset. A.** cryoSPARC processing workflow for obtaining consensus, local refined, and composite maps of the expanded state. **B.** Representative micrograph and **C.** 2D class average images.

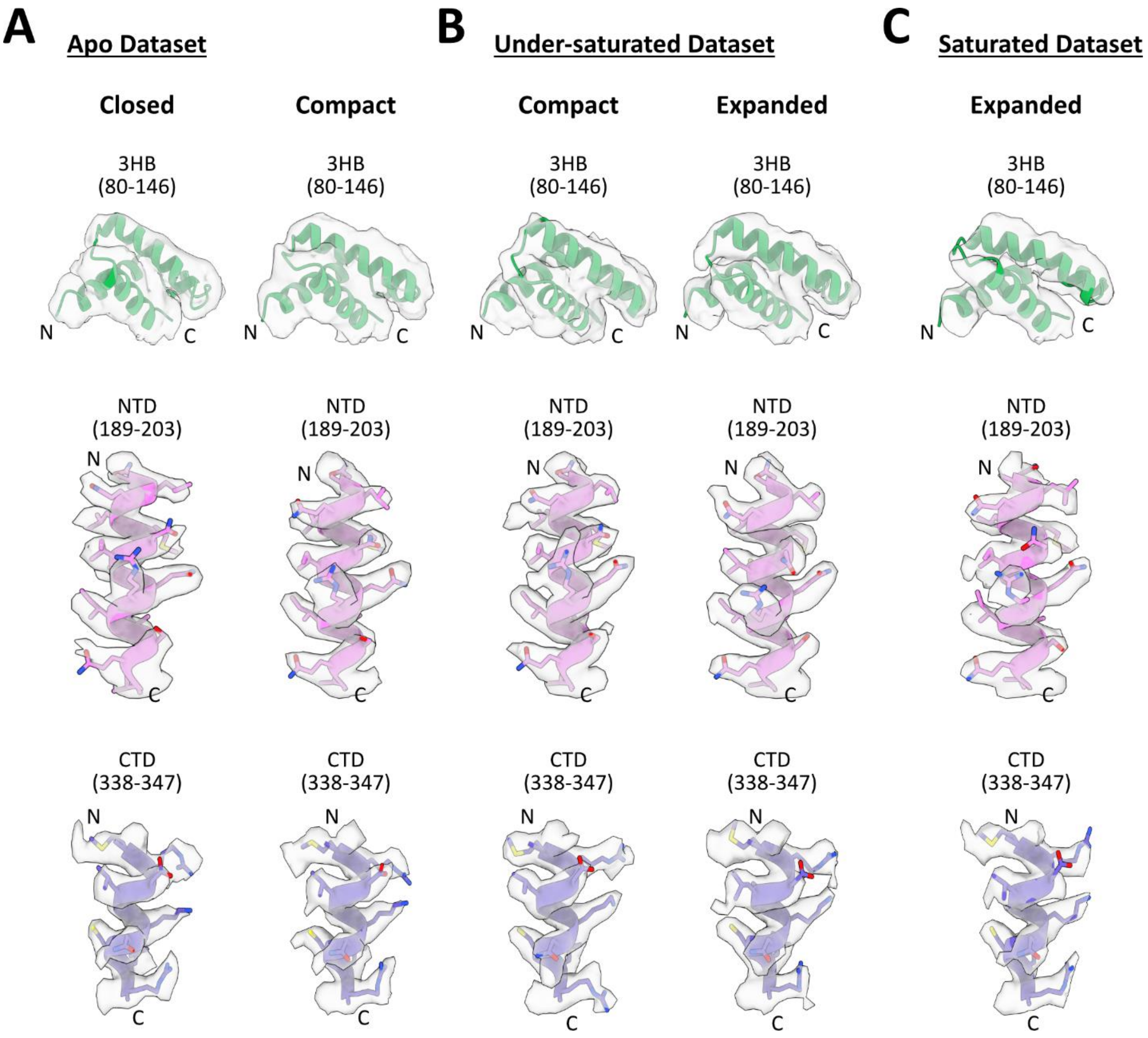


**Supplementary Figure 7. Examples of model-in-map fit quality. A-C.** (top) Fit of the atomic models into the respective local refined maps of 3HB between residues D80 and L146. Side chains in this region were not built due to lower map resolution. Fit of the atomic models into the consensus maps of: (middle) the NTD between residues N189 and V203; and (bottom) the CTD between residues M338 and R347.

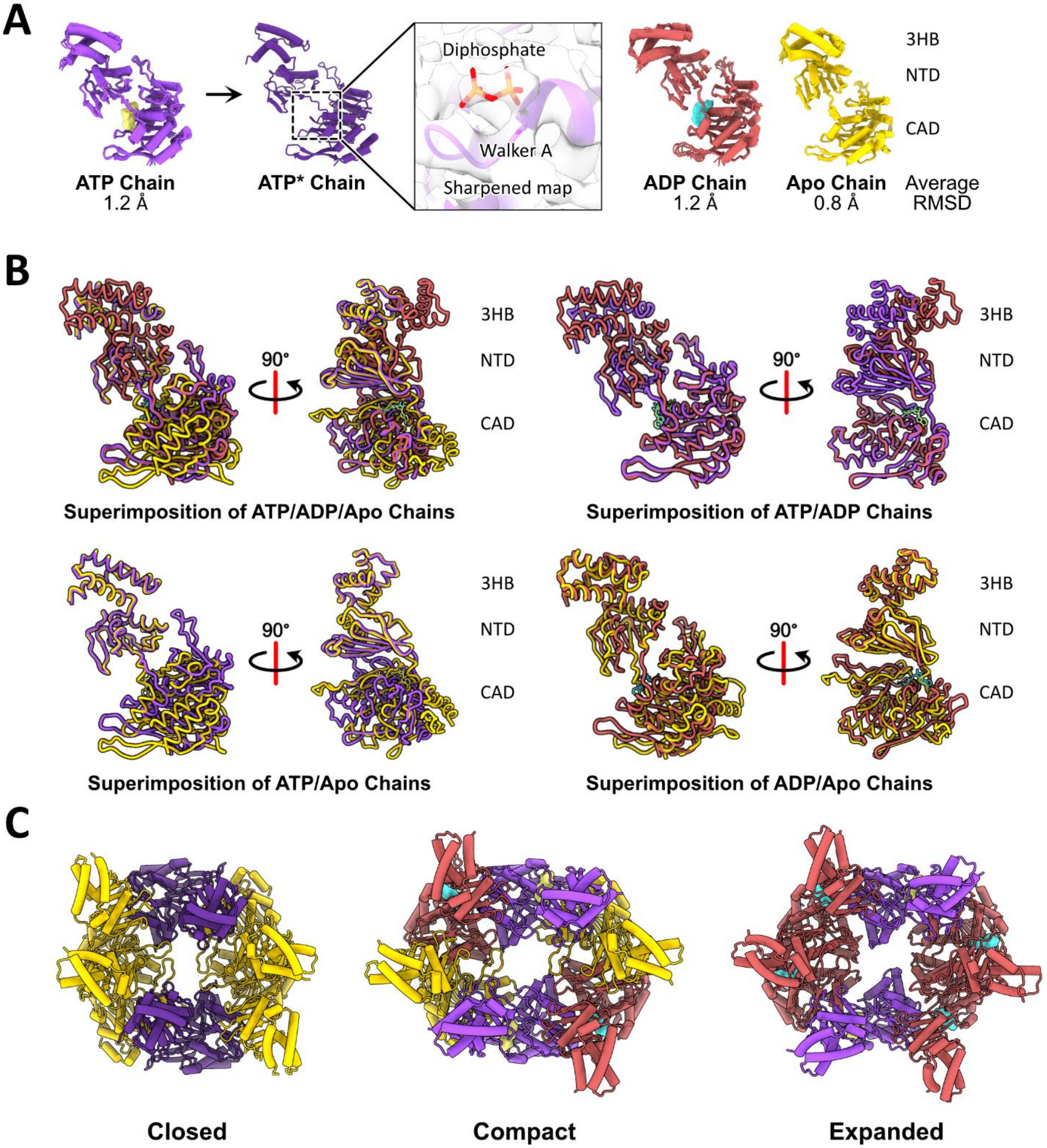


**Supplementary Figure 8. Classification and structural alignment of monomeric chains by nucleotide identity. A.** Superimposition of Cα atoms from all eighteen chains of the three structures grouped by their bound nucleotide. The average RMSD within each group is shown. Two chains within the ATP group, originating from the closed structure, adopt an ATP-like conformation despite the absence of ATP in the active site, which were designated ATP*. (inset) The residual density present in only the sharpened consensus map of the closed state could be fit with a diphosphate. **B.** Pairwise alignment of the ATP, ADP, and Apo chains from the compact structure, as well as alignment of all three chains, highlighting nucleotide-induced domain movements. Chains are colored as in panel A. **C.** Atomic models of the closed, compact, and expanded hexamers, colored according to the nucleotide bound in each chain as depicted in panel A.

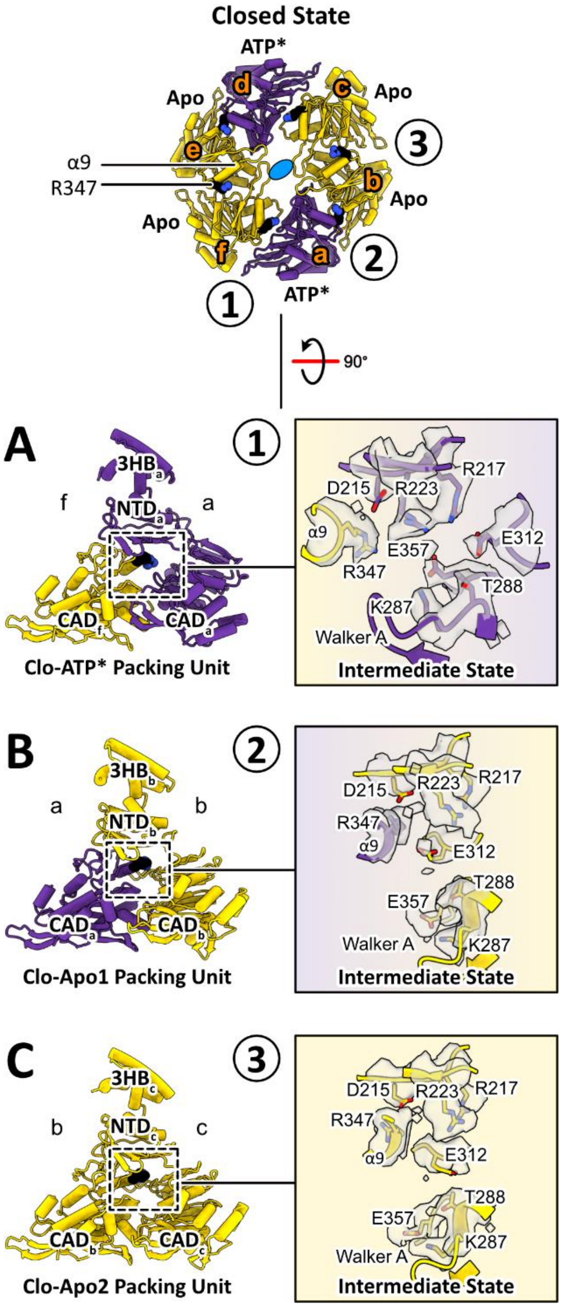


**Supplementary Figure 9. Packing unit analysis of the closed structure. A-C.** Packing units are numbered 1 to 3 according to their positions within an asymmetric unit of the hexamer. Full chains and their constituent domains are labelled from a to f and colored according to their bound nucleotides – ATP* in dark purple and Apo in gold. The residue R347 on the α9 helix in black is shown as a space-filling model colored by its side chain heteroatom. Active site analysis highlights residues, shown in stick representation, involved in stabilizing bound nucleotides. All residues and nucleotides are encased in their corresponding cryo-EM density (grey for amino acids) contoured at the same threshold level. The side chain rotamers of R217, R223, and R347 are classified as "intermediate".

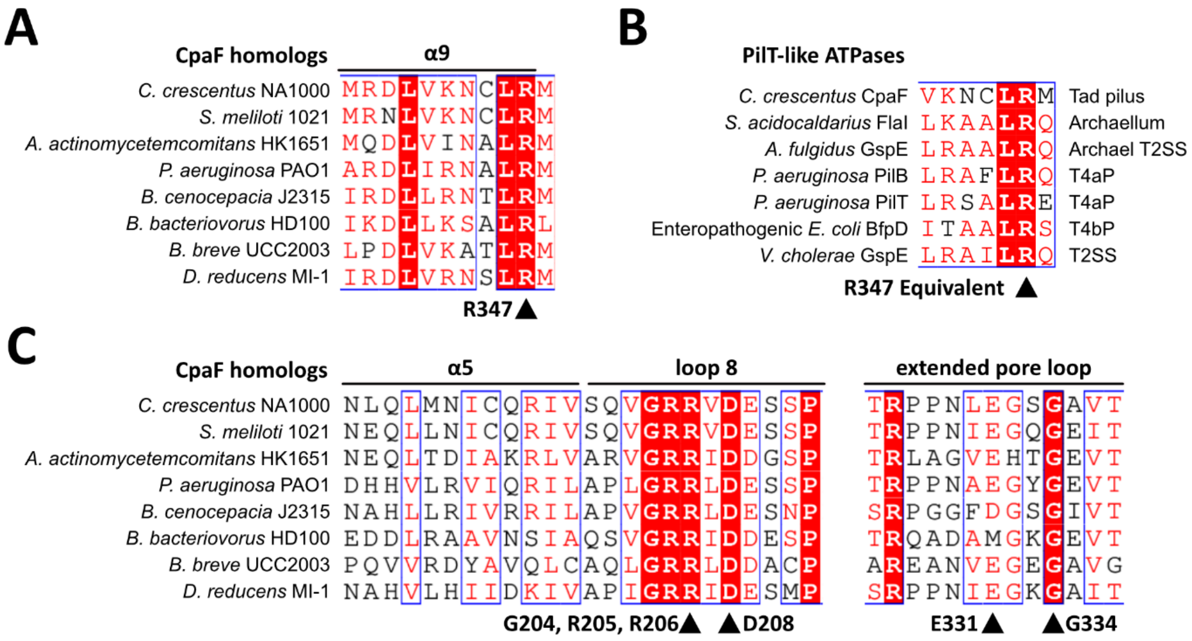


**Supplementary Figure 10. Multiple sequence alignments of CpaF.** Multiple sequence alignment of **(A)** representative CpaF orthologs from all major bacterial phyla, depicting conserved residue R347 on α9 helix, and (**B)** of PilT-like ATPases from all TFF superfamily systems, highlighting conservation of the R347 residue or its equivalent. (**C**) Multiple sequence alignment of representative CpaF orthologs from all major bacterial phyla, highlighting conserved residues in loop 8 and the extended pore loop.

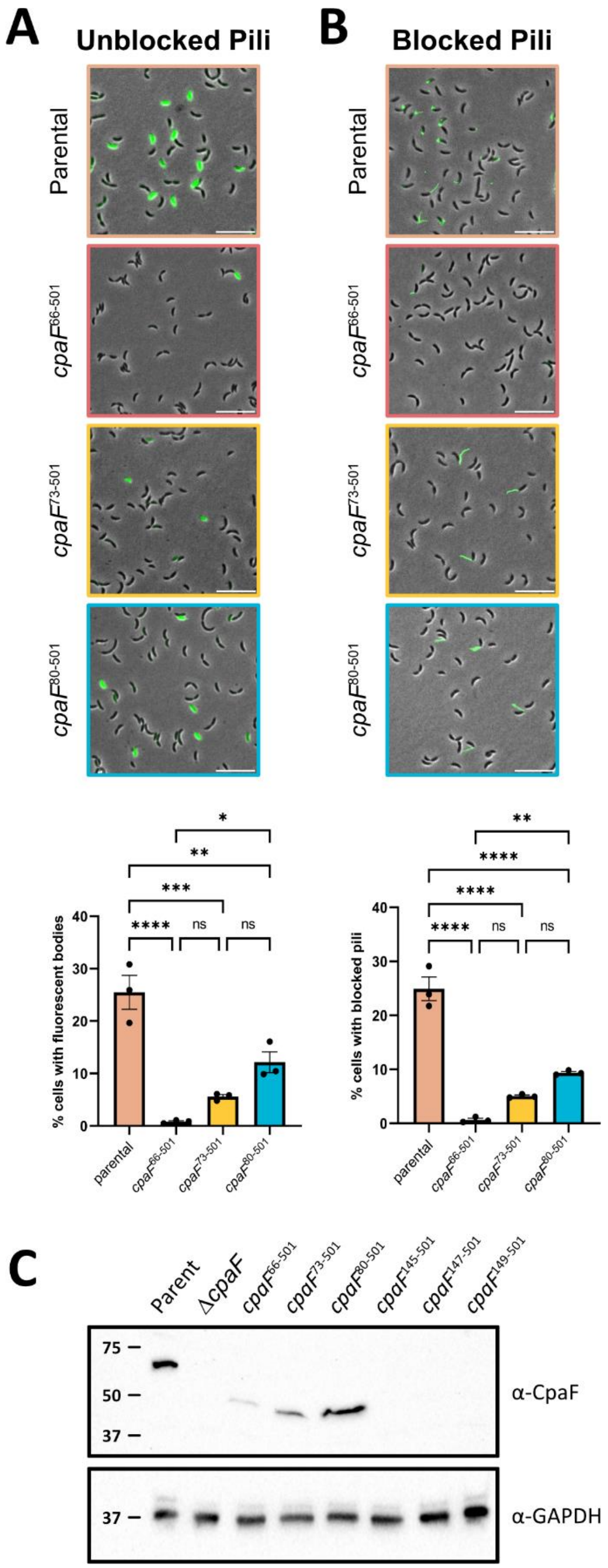


**Supplementary Figure 11. IDR truncations of CpaF at the native chromosomal locus. A.** (top) Representative images of a mixed population of *C. crescentus pil*-cys (parental) cells harboring the indicated IDR truncations of *cpaF* at the native chromosomal locus labeled with AF488-

maleimide. (bottom) Quantification of cell body fluorescence. **B.** (top) Representative images of a mixed population of *C. crescentus pil*-cys (parental) cells harboring the indicated IDR truncations of *cpaF* at the native locus, blocked with PEG5000-maleimide and labeled with AF488-maleimide. (bottom) Quantification of cells with blocked pili. In panels A and B, the scale bars in the microscopy images equal 10 µm, and each data point represents one biological replicate with three technical replicates. Statistics were determined using Tukey's multiple comparisons test. *$p < 0.05$. **$p < 0.01$. ***$p < 0.001$. ****$p < 0.0001$. ns, not significant. Error bars show standard error of the mean (SEM). **C.** Western blot showing expression of CpaF IDR and 3HB truncations from whole cell lysates probed using CpaF-specific antibodies. Antibody against GAPDH was used as a loading control.

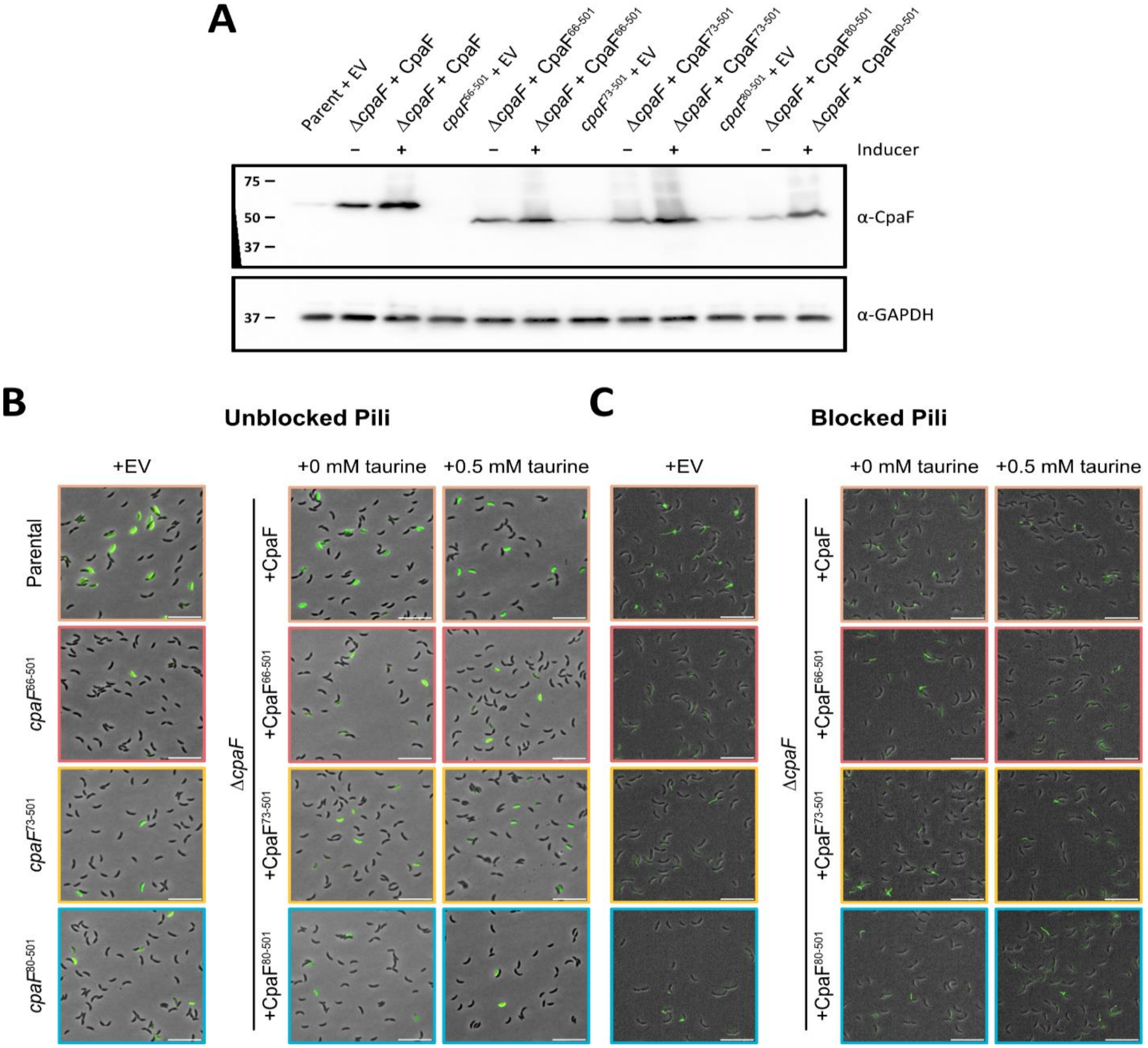


**Supplementary Figure 12. Overexpression of CpaF IDR truncation mutants produces similar pilus activity levels. A.** Western blot showing expression of CpaF IDR truncation constructs from whole cell lysates probed using CpaF-specific antibodies. Antibody against GAPDH was used as a loading control. **B.** Representative images of a mixed population of *C. crescentus pil*-cys (parental) cells harboring IDR truncations at the native chromosomal locus (left) or expressed from an inducible vector in a Δ*cpaF* background (right), labeled with AF488-maleimide. **C.** Representative images of a mixed population of *C. crescentus pil*-cys (parental) cells harboring IDR truncations at the native chromosomal locus (left) or expressed from an inducible vector in a Δ*cpaF* background, blocked with PEG5000-maleimide and labeled with AF488-maleimide. In panels A-C, cells harboring *cpaF* at the native chromosomal locus were transformed with empty vector (EV) as a control. Cells in a Δ*cpaF* background were transformed with an inducible vector harboring *cpaF* or truncation variants. Induction of *cpaF* expression was done via addition of 0.5 mM taurine. Scale bars = 10 μm.

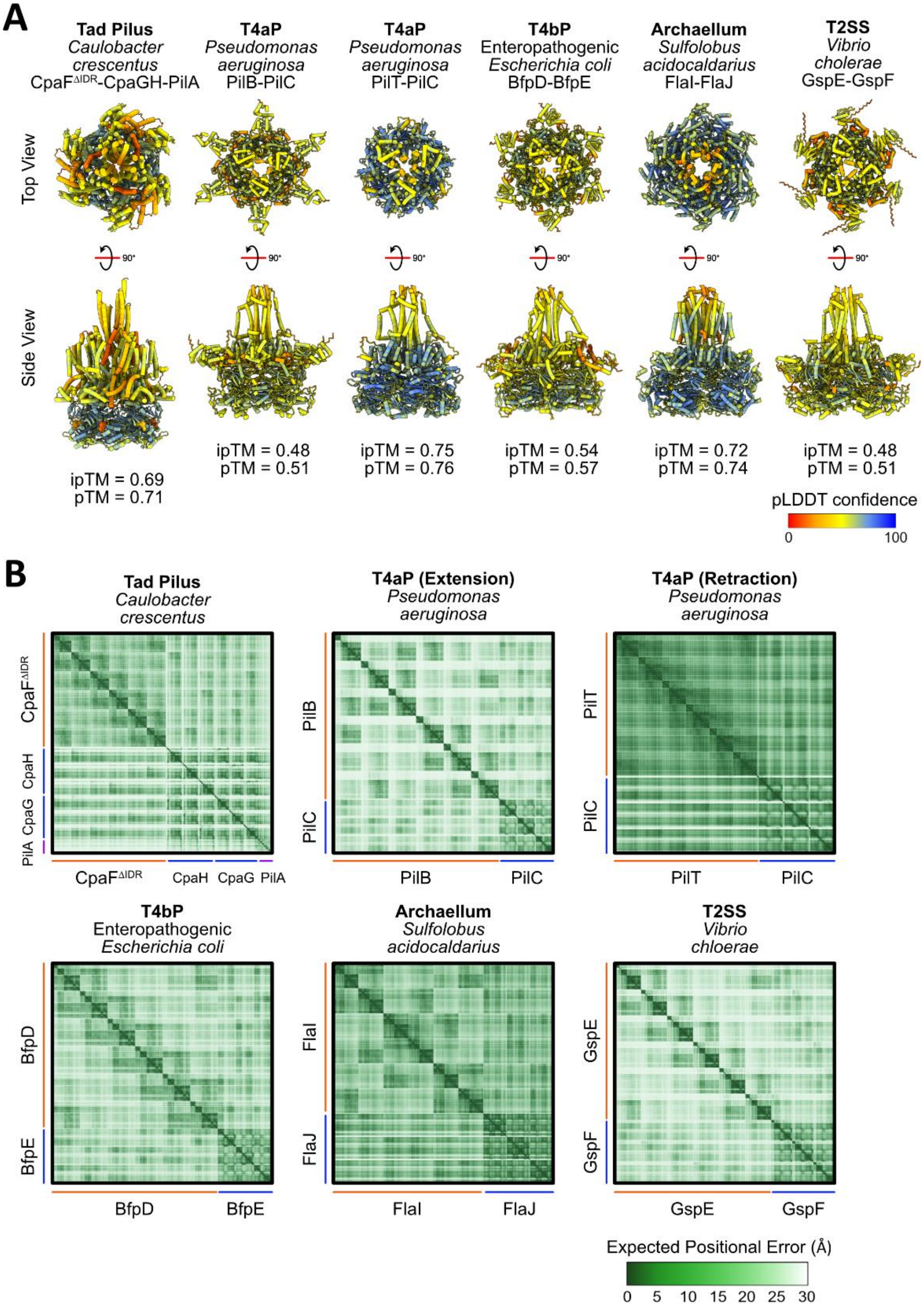


**Supplementary Figure 13. Alphafold3 prediction confidence of TFF superfamily motor subcomplexes. A.** Highest-ranking Alphafold3 predicted models of the motor subcomplexes of the TFF superfamily systems depicted in top and side views and colored by their respective predicted local distance difference test (pLDDT) scores, ranging from 0-100. The interface predicted template modeling (ipTM) and predicted template modeling (pTM) values from each prediction are shown at the bottom. **B.** Residue-residue predicted alignment error (PAE) plot for each prediction, which gives a distance error for every residue pair and estimates the positional error at a specific residue when the predicted and true structures are aligned. The values range from 0-30 Å. Corresponding proteins associated with each plot is shown with ATPases in orange, platform proteins in blue, and major pilin in purple.

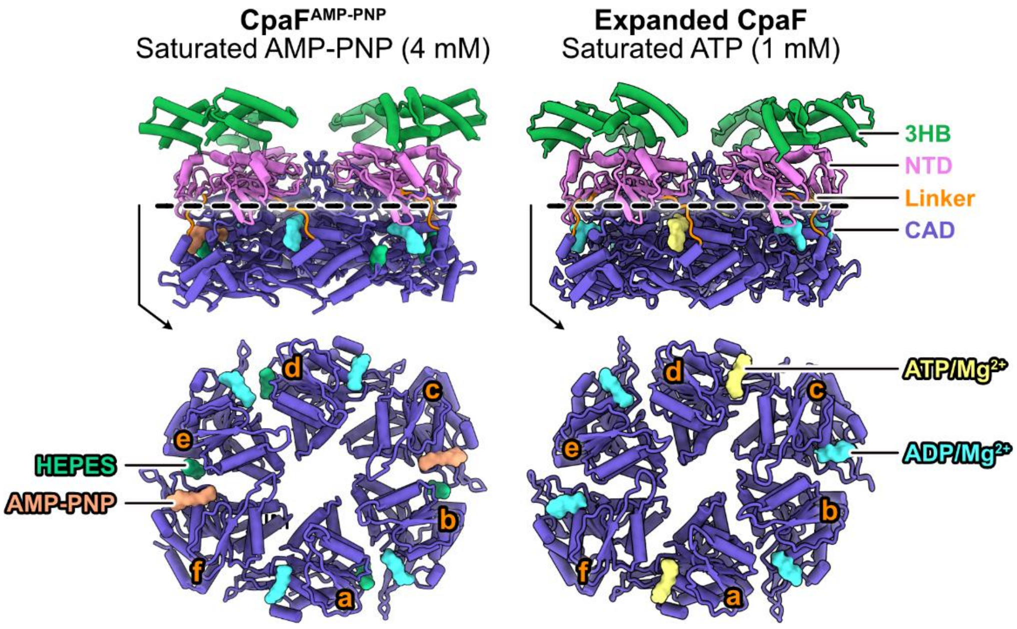


**Supplementary Figure 14. Comparison of nucleotide occupancy of two independently determined CpaF structures.** (Left) Atomic model of $\mathrm{CpaF_{AMP\text{-}PNP}}$ (PDB: 8RKD) shown from the side view and as a cross-section through the NTD (omitted) and CAD (shown). 2 AMP-PNP and 4 ADP molecules occupy the six active sites, while four additional HEPES molecules are bound between chains. (Right) Atomic model of the expanded state (this study) shown in the same orientations as $\mathrm{CpaF_{AMP\text{-}PNP}}$. 2 ATP and 4 ADP molecules occupy the six active sites.

### Supplementary Table 1. Cryo-EM collection, refinement, validation statistics

| | Closed – Apo (EMD-47431) (PDB 9E24) | Compact - Apo (EMD-47434) (PDB 9E25) | Compact - Under-saturated ATP/ADP (EMD-47437) (PDB 9E26) | Expanded - Under-saturated ATP/ADP (EMD-47440) (PDB 9E27) | Expanded - Saturated ATP (EMD-47444) (PDB 9E29) |
|---|---|---|---|---|---|
| **Data Collection and Processing** | | | | | |
| Magnification | 75,000x | 75,000x | 75,000x | 75,000x | 75,000x |
| Voltage (kV) | 300 | 300 | 300 | 300 | 300 |
| Electron Exposure ($e^{-}$/Å$^{2}$) | 36.7<br>42 | 36.7<br>42 | 36.7 | 36.7 | 42 |
| Tilt Angle | 0° | 0° | 0°<br>30° | 0°<br>30° | 0° |
| Defocus range (µm) | 1.0-2.0 | 1.0-2.0 | 1.0-2.0 | 1.0-2.0 | 1.0-2.0 |
| Pixel Size (Å) | 1.03 | 1.03 | 1.03 | 1.03 | 1.03 |
| Symmetry Imposed | C2 | C2 | C2 | C2 | C2 |
| Initial Particle Images (no.) | 1,078,013<br>1,716,591 | 1,078,013<br>1,716,591 | 1,605,149<br>473,550 | 1,605,149<br>473,550 | 2,250,189 |
| Final Particle Images (no.) | 84,636 | 149,839 | 226,031 | 138,569 | 273,260 |
| Map Resolution (Å) | 3.4 | 3.2 | 2.8 | 3.2 | 3.3 |
| FSC Threshold | 0.143 | 0.143 | 0.143 | 0.143 | 0.143 |
| Map Resolution Range (Å) | 2.2-10.1 | 2.2-9.3 | 2.2-26.7 | 2.2-9.6 | 2.2-9.4 |
| **Refinement** | | | | | |
| Initial Model Used | Alphafold2 | Alphafold2 | Alphafold2 | Alphafold2 | Alphafold2 |
| Model Resolution (Å) | 3.6 | 3.4 | 3.0 | 3.5 | 3.5 |
| FSC Threshold | 0.5 | 0.5 | 0.5 | 0.5 | 0.5 |
| Sharpening *B* Factor (Å) | Sharpened Locally | Sharpened Locally | Sharpened Locally | Sharpened Locally | Sharpened Locally |
| Model Composition | | | | | |
| Non-Hydrogen Atoms | 35898 | 35473 | 36064 | 36138 | 36141 |
| Protein Residues | 2532 | 2532 | 2532 | 2532 | 2532 |
| Ligands | 0 | 0 | 8 | 12 | 12 |
| Mean *B* factors (Å$^{2}$) | | | | | |
| Protein | 147.38 | 130.62 | 123.39 | 159.77 | 171.68 |
| Ligand | 0 | 0 | 111.05 | 151.98 | 149.59 |
| R.m.s Deviations | | | | | |
| Bond Lengths (Å) | 0.003 | 0.003 | 0.005 | 0.003 | 0.004 |
| Bond Angles (°) | 0.496 | 0.461 | 0.536 | 0.524 | 0.593 |
| Validation | | | | | |
| MolProbity Score | 1.48 | 1.57 | 1.50 | 1.75 | 1.88 |
| Clashscore | 6.55 | 5.24 | 5.82 | 8.52 | 10.21 |
| Poor Rotamers (%) | 0.06 | 0.06 | 1.07 | 0.11 | 0.17 |
| Ramachandran plot | | | | | |
| Favored (%) | 97.42 | 95.87 | 97.14 | 95.79 | 94.96 |
| Allowed (%) | 2.58 | 4.13 | 2.86 | 4.21 | 5.04 |
| Disallowed (%) | 0 | 0 | 0 | 0 | 0 |

| | Local Refined Closed – Apo (EMD-47432) | Local Refined Compact - Apo (EMD-47435) | Local Refined Compact - Under-saturated ATP/ADP (EMD-47438) | Local Refined Expanded - Under-saturated ATP/ADP (EMD-47441) | Local Refined Expanded - Saturated ATP (EMD-47445) |
|---|---|---|---|---|---|
| **Data Collection and Processing** | | | | | |
| Magnification | 75,000x | 75,000x | 75,000x | 75,000x | 75,000x |
| Voltage (kV) | 300 | 300 | 300 | 300 | 300 |
| Electron Exposure ($e^{-}$/Å$^{2}$) | 36.7<br>42 | 36.7<br>42 | 36.7 | 36.7 | 42 |
| Tilt Angle | 0° | 0° | 0°<br>30° | 0°<br>30° | 0° |
| Defocus range (µm) | 1.0-2.0 | 1.0-2.0 | 1.0-2.0 | 1.0-2.0 | 1.0-2.0 |
| Pixel Size (Å) | 1.03 | 1.03 | 1.03 | 1.03 | 1.03 |
| Initial Particle Images (no.) | 1,078,013<br>1,716,591 | 1,078,013<br>1,716,591 | 1,605,149<br>473,550 | 1,605,149<br>473,550 | 2,250,189 |
| Final Particle Images (no.) C2 symmetry expanded | 169,272 | 299,678 | 452,062 | 277,138 | 546,520 |
| Map Resolution (Å) | 4.0 | 4.1 | 3.2 | 3.6 | 3.7 |
| FSC Threshold | 0.143 | 0.143 | 0.143 | 0.143 | 0.143 |
| Map Resolution Range (Å) | 2.5-50.2 | 2.4-46.9 | 2.2-39.4 | 2.2-47.4 | 2.3-41.5 |

**Supplementary Table 2. Strains and CpaF ortholog accession numbers used for phylogenetic analysis in Fig. 4.**

See Excel spread sheet.

**Supplementary Table 3. Bacterial/archaeal species, proteins, and their associated UniProt accession numbers used for Alphafold3 predictions in Fig. 7.**

| System | Organism | ATPase | | Platform | |
|---|---|---|---|---|---|
| | | Protein | Accession # | Protein | Accession # |
| T4aP | *Pseudomonas aeruginosa* | PilB | P22608 | PilC | P22609 |
| T4aP | *Pseudomonas aeruginosa* | PilT | P24559 | PilC | P22609 |
| T4bP | Enteropathogenic *Escherichia coli* | BfpD | B7UTD6 | BfpE | B7UTD7 |
| Tad Pilus | *Caulobacter crescentus* | CpaF | A0A0H3CDS2 | CpaG<br>CpaH | A0A0H3CC65<br>A0A0H3CC31 |
| Archaellum | *Sulfolobus acidocaldarius* | FlaI | Q4J9L0 | FlaJ | Q4J9L1 |
| T2SS | *Vibrio cholerae* | GspE | A0A7Z7VKT0 | GspF | A0A7Z7VJS7 |

**Supplementary Table 4. Bacterial strains, plasmids, and primers used in this study.**

| Strain | Description | Source |
|---|---|---|
| ***Escherichia coli* strains** | | |
| DH5α | Cloning strain; $F^{-}$*mcrA* Δ(*mrr-hsd*RMS-*mcr*BC) φ80*lac*ZΔM15 Δ*lac*X74 *rec*A1 *ara*D139 Δ(*ara-leu*)7697 *gal*U *gal*K λ–*rps*L($Str^{R}$) *end*A1 *nup*G | Invitrogen |
| Rosetta 2 (DE3) | Protein expression strain; $F^{-}$ *ompT hsdS*$_B$($r_B^{-}$ $m_B^{-}$) *gal dcm* (DE3) pRARE2 ($Cam^{R}$) | Stratagene |
| NEB5α | DH5α derivative, *fhuA2Δ*(*argF-lacZ*)*U169 phoA glnV44 Φ80Δ*(*lacZ*)*M15 gyrA96 recA1 relA1 endA1 thi-1 hsdR17* | New England Biolabs |
| YB8445 | NEB5α pNPTS138::Δ*cpaF* | This study |
| YB10220 | NEB5α pNPTS138::$cpaF^{66-501}$ | This study |
| YB10221 | NEB5α pNPTS138::$cpaF^{73-501}$ | This study |
| YB10222 | NEB5α pNPTS138::$cpaF^{80-501}$ | This study |
| YB10223 | NEB5α pNPTS138::$cpaF^{145-501}$ | This study |
| YB10224 | NEB5α pNPTS138::$cpaF^{147-501}$ | This study |
| YB10225 | NEB5α pNPTS138::$cpaF^{149-501}$ | This study |
| YB10226 | NEB5α $pJC585^{-}$ | This study |
| YB10227 | NEB5α pJC585::*cpaF* | This study |
| YB10228 | NEB5α pJC585::$cpaF^{66-501}$ | This study |
| YB10229 | NEB5α pJC585::$cpaF^{73-501}$ | This study |
| YB10230 | NEB5α pJC585::$cpaF^{80-501}$ | This study |
| ***Caulobacter crescentus* strains** | | |
| NA1000 | Synchronizable *C. crescentus* lab adapted strain that does not produce holdfast | [56] |
| YB8288 | NA1000 $pilA^{T36C}$, pili can be labelled with maleimide-conjugated fluorophores | [5] |
| YB8446 | NA1000 $pilA^{T36C}$ Δ*cpaF*, allelic exchange with plasmid from YB8445 electroporated into YB8288 | This study |
| YB10231 | NA1000 $pilA^{T36C}$ $cpaF^{66-501}$, allelic exchange with plasmid from YB10220 electroporated into YB8288 | This study |
| YB10232 | NA1000 $pilA^{T36C}$ $cpaF^{73-501}$, allelic exchange with plasmid from YB10221 electroporated into YB8288 | This study |
| YB10233 | NA1000 $pilA^{T36C}$ $cpaF^{80-501}$, allelic exchange with plasmid from YB10222 electroporated into YB8288 | This study |
| YB10234 | NA1000 $pilA^{T36C}$ $cpaF^{145-501}$, allelic exchange with plasmid from YB10223 electroporated into YB8288 | This study |
| YB10235 | NA1000 $pilA^{T36C}$ $cpaF^{147-501}$, allelic exchange with plasmid from YB10224 electroporated into YB8288 | This study |
| YB10236 | NA1000 $pilA^{T36C}$ $cpaF^{149-501}$, allelic exchange with plasmid from YB10225 electroporated into YB8288 | This study |

| | | |
|---|---|---|
| YB10237 | NA1000 *pilA*$^{T36C}$ pJC585$^{-}$, electroporation of plasmid from YB10226 into YB8288 | This study |
| YB10238 | NA1000 *pilA*$^{T36C}$ *cpaF*$^{66\text{-}501}$ pJC585$^{-}$, electroporation of plasmid from YB10226 into YB10231 | This study |
| YB10239 | NA1000 *pilA*$^{T36C}$ *cpaF*$^{73\text{-}501}$ pJC585$^{-}$, electroporation of plasmid from YB10226 into YB10232 | This study |
| YB10240 | NA1000 *pilA*$^{T36C}$ *cpaF*$^{80\text{-}501}$ pJC585$^{-}$, electroporation of plasmid from YB10226 into YB10233 | This study |
| YB10241 | NA1000 *pilA*$^{T36C}$ Δ*cpaF* pJC585::*cpaF*, electroporation of plasmid from YB10227 into YB8466 | This study |
| YB10242 | NA1000 *pilA*$^{T36C}$ Δ*cpaF* pJC585::*cpaF*$^{66\text{-}501}$, electroporation of plasmid from YB10228 into YB8466 | This study |
| YB10243 | NA1000 *pilA*$^{T36C}$ Δ*cpaF* pJC585::*cpaF*$^{73\text{-}501}$, electroporation of plasmid from YB10229 into YB8466 | This study |
| YB10244 | NA1000 *pilA*$^{T36C}$ Δ*cpaF* pJC585::*cpaF*$^{80\text{-}501}$, electroporation of plasmid from YB10230 into YB8466 | This study |
| **Plasmid** | | |
| pNPTS138 | Litmus 38 derivative, *nptI oriT sacB, Kan*$^{R}$; used for allelic exchange in *C. crescentus* | M.R.K Alley, unpublished |
| pNPTS138::Δ*cpaF* | pNPTS138 containing 646 bp upstream of *cpaF* codon 17 fused to 637 bp downstream of *cpaF* codon 486 at the EcoRV site; used to generate an in-frame, markerless deletion of the *cpaF* ORF from the NA1000 genome | This study |
| pNPTS138::*cpaF*$^{66\text{-}501}$ | pNPTS138 containing 477 bp upstream of and including the *cpaF* start codon fused to 514 bp downstream of *cpaF* codon 65 at the EcoRV site; used to remove codons 2-65 of the *cpaF* ORF from the NA1000 genome | This study |
| pNPTS138::*cpaF*$^{73\text{-}501}$ | pNPTS138 containing 477 bp upstream of and including the *cpaF* start codon fused to 496 bp downstream of *cpaF* codon 72 at the EcoRV site; used to remove codons 2-72 of the *cpaF* ORF from the NA1000 genome | This study |
| pNPTS138::*cpaF*$^{80\text{-}501}$ | pNPTS138 containing 477 bp upstream of and including the *cpaF* start codon fused to 472 bp downstream of *cpaF* codon 79 at the EcoRV site; used to remove codons 2-79 of the *cpaF* ORF from the NA1000 genome | This study |
| pNPTS138::*cpaF*$^{145\text{-}501}$ | pNPTS138 containing 477 bp upstream of and including the *cpaF* start codon fused to 506 bp downstream of *cpaF* codon 144 at the EcoRV site; used to remove codons 2-144 of the *cpaF* ORF from the NA1000 genome | This study |
| pNPTS138::*cpaF*$^{147\text{-}501}$ | pNPTS138 containing 477 bp upstream of and including the *cpaF* start codon fused to 500 bp downstream of *cpaF* codon 146 at the EcoRV site; used to remove codons 2-146 of the *cpaF* ORF from the NA1000 genome | This study |

| | | |
|---|---|---|
| pNPTS138::*cpaF*$^{149-501}$ | pNPTS138 containing 477 bp upstream of and including the *cpaF* start codon fused to 494 bp downstream of *cpaF* codon 148 at the EcoRV site; used to remove codons 2-148 of the *cpaF* ORF from the NA1000 genome | This study |
| pJC585 | *nptII tauR Ptau-rfp, Kan*$^R$; RFP reporter under the control of a taurine inducible promoter | J. Chen, unpublished |
| pJC585$^-$ | pJC585 digested with KpnI to remove part of the vector-encoded rfp sequence, used as an empty vector control | This study |
| pJC585::*cpaF* | pJC585 containing *cpaF* fused to a synthetic RBS, inserted between the EcoRI and BamHI sites, under the control of a taurine-inducible promoter | This study |
| pJC585::*cpaF*$^{66-501}$ | pJC585 containing *cpaF*$^{66-501}$ fused to a synthetic RBS, inserted between the EcoRI and BamHI sites, under the control of a taurine-inducible promoter | This study |
| pJC585::*cpaF*$^{73-501}$ | pJC585 containing *cpaF*$^{73-501}$ fused to a synthetic RBS, inserted between the EcoRI and BamHI sites, under the control of a taurine-inducible promoter | This study |
| pJC585::*cpaF*$^{80-501}$ | pJC585 containing *cpaF*$^{80-501}$ fused to a synthetic RBS, inserted between the EcoRI and BamHI sites, under the control of a taurine-inducible promoter | This study |
| pET28a | IPTG-inducible expression vector encoding N-terminal hexa-histidine tag, a thrombin cleavage site, and an optional C-terminal hexahistidine tag, Kan$^R$ | Novagen |
| pET28a::CpaF | pET28a with WT *C. crescentus* NA1000 *cpaF* fused to an N-terminal hexa-histidine tag; Kan$^R$ | This study |
| pET28a::CpaF$^{66-501}$ | pET28a::CpaF corresponding to residues 66-501 | This study |
| pET28a::CpaF$^{73-501}$ | pET28a::CpaF corresponding to residues 73-501 | |
| pET28a::CpaF$^{80-501}$ | pET28a::CpaF corresponding to residues 80-501 | This study |
| **Primer** | | |
| pET28cpaF-FI | **GCGCGGCAGCCAT**ATGTTCGGAAAGCGCGACTCGTCAGC | This study |
| pET28cpaF-RI | **CTTGTCGACGGAGCTCGAATTC**CTACTCCGCCGCGTCGAGGGCTT | This study |
| pET28cpaF-FV | AAGCCCTCGACGCGGCGGAGTAG**GAATTCGAGCTCCGTCGACAAG** | This study |
| pET28cpaF-RV | GCTGACGAGTCGCGCTTTCCGAACAT**CATATGGCTGCCGCGC** | This study |
| pET28cpaF66-FI | **GCCGCGCGGCAGCCAT**CAGGGCCAGCCGCAAACGG | This study |
| pET28cpaF66-RI | **CCGCAAGCTTGTCGACGGAGCTCGAATTC**CTACTCCGCCGCGTCGAGGGCTT | This study |
| pET28cpaF66-FV | **AAGCCCTCGACGCGGCGGAGTAG**GAATTCGAGCTCCGTCGACAAGCTTGCGG | This study |

| | | |
|---|---|---|
| pET28cpaF66-RV | CCGTTTGCGGCTGGCCCTG**ATGGCTGCCGCGCGGC** | This study |
| pET28cpaF73-FI | **GCCGCGCGGCAGCCAT**AACATCGTCCGTGAGCAGAGCGACTAC | This study |
| pET28cpaF73-RI | **CCGCAAGCTTGTCGACGGAGCTCGAATTC**CTACTCCGCCGCGTCGAGGGCTT | This study |
| pET28cpaF73-FV | AAGCCCTCGACGCGGCGGAGTAG**GAATTCGAGCTCCGTCGACAAGCTTGCGG** | This study |
| pET28cpaF73-RV | GTAGTCGCTCTGCTCACGGACGATGTT**ATGGCTGCCGCGCGGC** | This study |
| pET28cpaF80-FI | **GCCGCGCGGCAGCCAT**GACTACTACCACGCCACCAAGACCACGATCTTC | This study |
| pET28cpaF80-RI | **CCGCAAGCTTGTCGACGGAGCTCGAATTC**CTACTCCGCCGCGTCGAGGGCTT | This study |
| pET28cpaF80-FV | **AAGCCCTCGACGCGGCGGAGTAG**GAATTCGAGCTCCGTCGACAAGCTTGCGG | This study |
| pET28cpaF80-RV | **GAAGATCGTGGTCTTGGTGGCGTGGTAGTAGTC**ATGGCTGCCGCGCGGC | This study |
| Δ*cpaF*-upF | **AATTCTGGAT**CCAGCCTGACCGCCTG | This study |
| Δ*cpaF*-upR | *AGCCCGTA*CGGCGCCTTGGGATCGCC | This study |
| Δ*cpaF*-downF | *AGGCGCCG*TACGGGCTGGAGCGCGAGC | This study |
| Δ*cpaF*-downR | **AAGCTTCCTGCAG**TCTCGGTCAGGAT | This study |
| *cpaF*-trunc-upF | **GCCAAGCTTCTCTGCAGGAT**TTCGAGGAAGTGACCCAGAAGATCC | This study |
| *cpaF*-trunc-66-upR | *CCGTTTGCGGCTGGCCCTG*CATCTACTTCTTCTTGAACAGGCCCGAG | This study |
| *cpaF*-trunc-66-downF | CAGGGCCAGCCGCAAACGG | This study |
| *cpaF*-trunc-73-upR | *TAGTCGCTCTGCTCACGGACGATGTT*CATCTACTTCTTCTTGAACAGGCCCGAG | This study |
| *cpaF*-trunc-73-downF | AACATCGTCCGTGAGCAGAGCGACTA | This study |
| *cpaF*-trunc-80-upR | *GGTCTTGGTGGCGTGGTAGTAGTC*CATCTACTTCTTCTTGAACAGGCCCGAG | This study |
| *cpaF*-trunc-80-downF | GACTACTACCACGCCACCAAGACC | This study |
| *cpaF*-trunc-145-upR | *AGCGGGCCATAGCCGAGGAC*CATCTACTTCTTCTTGAACAGGCCCGAG | This study |
| *cpaF*-trunc-145-downF | GTCCTCGGCTATGGCCCGCT | This study |
| *cpaF*-trunc-147-upR | *GCTCCAGCGGGCCATAGCC*CATCTACTTCTTCTTGAACAGGCCCGAG | This study |

| | | |
|---|---|---|
| *cpaF*-trunc-147-downF | GGCTATGGCCCGCTGGAGC | This study |
| *cpaF*-trunc-149-upR | *CAGCGGCTCCAGCGGGCC*CATCTACTTCTTCTTGAACAGGCCCGAG | This study |
| *cpaF*-trunc-149-downF | GGCCCGCTGGAGCCGCTG | This study |
| *cpaF*-trunc-IDR-downR | **GCGAATTCGTGGATCCAGAT**TCGGACCATCCAGCGCCAAGG | This study |
| *cpaF*-trunc-3HB-downR | **GCGAATTCGTGGATCCAGAT**AGTTCGGCGGCGTCCTCGCA | This study |
| *cpaF*-comp-F | GTGTG**GAATTC*TTTAAGAAGGAGATATACAT***ATGTTCGGAAAGCGCGACTCGTCAG | This study |
| *cpaF*-comp-66-F | GAGTG**GAATTC*TTTAAGAAGGAGATATACAT***ATGCAGGGCCAGCCGCAAACGG | This study |
| *cpaF*-comp-73-F | GTGT**GGAA*TTCTTTAAGAAGGAGATATACAT***ATGAACATCGTCCGTGAGCAGAGCGACTA | This study |
| *cpaF*-comp-79-F | GTGT**GGAA*TTCTTTAAGAAGGAGATATACAT***ATGGACTACTACCACGCCACCAAGACC | This study |
| *cpaF*-comp-R | GTGT**GGGATCC**CTACTCCGCCGCGTCGAGG | This study |

*Restriction sites and sequences for Gibson assembly into destination plasmids are bolded; regions of complementarity to the target amplicon are underlined; regions of reverse complementarity (to facilitate splicing) are italicized; synthetic ribosomal binding sites are in bold italics.

**Supplementary Movie 1.** Interpolated trajectory between the Com-Apo and Exp-ATP packing units during an ATP binding event. The packing unit orientation corresponds to the views depicted in Fig. 3B.

**Supplementary Movie 2.** Interpolated trajectory between the Com-ATP and Exp-ADP1 packing units from an ATP hydrolysis event. The packing unit orientation corresponds to the views depicted in Fig. 3E.

**Supplementary Movie 3.** Interpolated trajectory between the Exp-ADP2 and Com-Apo packing units from an ADP release event. The packing unit orientation corresponds to the views depicted in Fig. 3G.

**Supplementary Movie 4.** Proposed clockwise rotary mechanism of CpaF catalysis. Iterative interpolated trajectories between the compact and expanded states and back to the compact state, depicting a total of 180° clockwise rotation about the symmetry axis. Each chain is labeled from a to f. The video may be viewed as a loop to visualize a full 360° clockwise rotation.
